# HeartBioPortal2.0: new developments and updates for genetic ancestry and cardiometabolic quantitative traits in diverse human populations

**DOI:** 10.1101/2020.10.19.346445

**Authors:** Bohdan B. Khomtchouk, Kasra A. Vand, Christopher S. Nelson, Salvator Palmisano, Robert L. Grossman

## Abstract

Cardiovascular disease (CVD) is the leading cause of death worldwide for both genders and across most racial and ethnic groups. However, different races and ethnicities exhibit different rates of cardiovascular disease and its related cardiorenal and metabolic co-morbidities, suggesting differences in genetic predisposition and risk of onset, as well as socioeconomic and lifestyle factors (diet, exercise, etc.) that act upon an individual’s unique underlying genetic background. Here we present HeartBioPortal2.0, a major update to HeartBioPortal, the world’s largest CVD genetics data precision medicine platform for harmonized CVD-relevant genetic variants, which now enables search and analysis of human genetic information related to heart disease across ethnically diverse populations and cardiovascular/renal/metabolic quantitative traits pertinent to CVD pathophysiology. HeartBioPortal2.0 is structured as a cloud-based computing platform and knowledge portal that consolidates a multitude of CVD-relevant next-generation sequencing data modalities into a single powerful query and browsing interface between data and user via a user-friendly web application publicly available to the scientific research community. Since its initial release, HeartBioPortal2.0 has added new cardiovascular/renal/metabolic disease relevant gene expression data as well as genetic association data from numerous large-scale genome-wide association study (GWAS) consortiums such as CARDIoGRAMplusC4D, TOPMed, FinnGen, AFGen, MESA, MEGASTROKE, UK Biobank, CHARGE, Biobank Japan, MyCode, among other studies. In addition, HeartBioPortal2.0 now includes support for quantitative traits and ethnically diverse populations, allowing users to investigate the shared genetic architecture of any gene or its variants across the continuous cardiometabolic spectrum from health (e.g., blood pressure traits) to disease (hypertension), facilitating the understanding of CVD trait genetics that inform health-to-disease transitions and endophenotypes. Custom visualizations in the new and improved user interface (UI), including performance enhancements and new security features such as user authentication collectively re-imagine HeartBioPortal’s user experience and provide a data commons that co-locates data, storage and computing infrastructure in the context of studying the genetic basis behind the leading cause of global mortality.

**Database URL:** https://www.heartbioportal.com/

## 1 Introduction

The world’s aging population (>65 years old) is projected to triple by the year 2050, creating an impending rise in future demand for healthcare innovation within the cardiovascular disease market sector, since the most important determinant of cardiovascular health is a person’s age (1), which is why age is such a critical component of cardiovascular disease (CVD) etiology. Specifically, age is known to be the largest risk factor of CVD (2), as it lowers the threshold for cardiovascular disease manifestation (3). CVDs are comprised of a group of disorders of the heart and blood vessels and include coronary artery disease, cerebrovascular disease (stroke), heart failure and other conditions, including cardiorenal and cardiometabolic co-morbidities and complications such as chronic kidney disease and type II diabetes. Given the rise in the aging population, coupled with the continuous decline of sequencing costs, genetics will continue to occupy an increasingly important role in the clinic. Many cardiology practice guidelines already incorporate genetic data in recommendations for diagnosis and personalized clinical management (4). Already it is well-recognized by clinicians and researchers alike that the roadmap to future implementation of personalized, or precision, medicine in cardiovascular/renal/metabolic disease must take into account individual and subgroup variability in genetic architecture, environment, clinical measures, lifestyle, cost-effectiveness and treatment burden, e.g., in disorders with substantial phenotypic heterogeneity such as type II diabetes (5) or heart failure (6–7). A step one to this process involves making it computationally feasible to figure out what is currently known about any given gene of interest in any specific CVD phenotype using the latest publicly available datasets/publications, and providing the tools/infrastructure to do so, effectively leveraging for secondary analysis existing clinical and genetic datasets from federally funded NIH/NSF research grants to provide an easily searchable and accessible data commons for all. Such a computational framework and cardioinformatics initiative (8) would mark a significant advance in traversing not only the bench-to-bedside divide but also the informatics-to-medicine divide that still exists in contemporary biomedicine and biomedical data science. The vision of HeartBioPortal (9) has always been to bridge these gaps by heralding in a data-driven future for cardiovascular clinical patient care, effectively building the computational supply needed to address future demand at the population levels projected in 2050 and beyond.

The HeartBioPortal project originally began as a way to lower the barriers to complex biological discovery from existing clinical and genetic datasets, similar to the dynamic provided by cancer genomics resources such as cBioPortal (10–11) and the Genomic Data Commons (12–13). Resources such as the American Heart Association’s Precision Medicine Platform (14) or Broad’s Cardiovascular Disease Knowledge Portal (15) either specialize on a single data modality (e.g., GWAS) or simply do not provide intuitive visualization and analysis of large-scale CVD datasets, instead relying on users to pinpoint relevant datasets themselves and generate their own independent analyses on them, putting these resources out of reach from biologists with no programming or bioinformatics expertise. This trend can still very much be observed in recently launched emerging resources such as NHLBI’s BioData Catalyst (16) initiative that does not provide a simple interface for users who simply wish to perform gene or variant-level search across existing CVD genomic datasets, which is an unmet need adequately addressed in HeartBioPortal. In essence, HeartBioPortal effectively democratizes CVD genomics data, by enabling gene and variant-level search across various heart disease and stroke phenotypes for the broader research community. At the same time, HeartBioPortal is also directly useful to power users interested in seeing CVD genetics information at scale, given that the platform already bioinformatically preprocesses and distills down terabyte-scale publicly available genomic data across multiple heterogenous modalities into easy-to-interpret figures and charts, including tidy data downloads provided in simple .csv format. Essentially, initiatives such as NHLBI’s BioData Catalyst (BDC) and knowledge portals such as HeartBioPortal serve different needs and purposes-one (BDC) provides access to CVD datasets and workspaces that support large-scale genomic pipelines, the other (HeartBioPortal) provides an easy to use capability to search, analyze and visualize annotated CVD variants that have been uniformly processed and curated. As detailed on BioData Catalyst’s Zenodo launched on August 31^st^, 2020: “Over the last several decades, NHLBI has invested in creating a significant resource for research and development by supporting the creation of many observational, epidemiological, and longitudinal datasets related to heart, lung, blood and sleep phenotypes, with the aim of uncovering insights that may be leveraged toward novel therapeutic, interventional, or preventive strategies resulting in improved patient outcomes. New technologies and favorable cost trajectories have enabled detailed characterization of these study participants including whole genome sequencing (and other omics) and imaging on hundreds of thousands of participants. Together this data coupled with animal and cellular models increase opportunities for data-driven translational science.” HeartBioPortal provides a simple and intuitive way to search, analyze and visualize some of the most important genomics variants in this data.

## 2 New Developments and Updates

Since its initial debut in May 2019, HeartBioPortal2.0 has more than tripled its number of unique CVD phenotypes (from 12 to 37), increased its clinical genetic variant store count by three orders of magnitude (from 5,792 to 6,906,292), almost doubled the number of human genic and intergenic regions featured (from 23,606 to 44,988), added 54 diverse subpopulations across 27 new studies/cohorts, introduced 9,641 more genes from differential expression analyses of various new high quality CVD-relevant gene expression studies published between now and initial debut, and added 56 cardiometabolic quantitative traits such as lipids, electrolytes, ECG/MRI traits, etc. to facilitate complex exploratory data analyses such as those presented in Figure 1. Given that the CVD research community continues to rely heavily on publicly available GWAS data and summary statistics, we have focused a lot of our software engineering effort on developing custom JavaScript visualizations such as the data-intensive interactive Venn diagram in Figure 1 to help make sense of GWAS data at scale in a fast and responsive UI that drives biological insight. We have also made other substantial UI improvements such as a new p-value dropdown menu directly in the Genetic Association panel at the top of HeartBioPortal2.0’s gene pages to allow the user to visually filter displayed GWAS results according to desired levels of statistical stringency.

**Figure 1.**
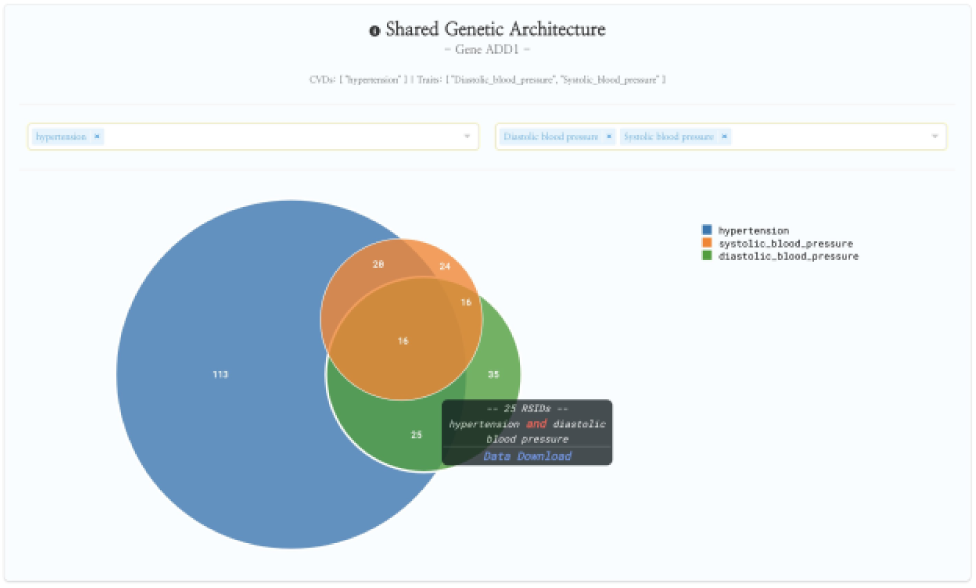
HeartBioPortal2.0’s shared genetic architecture feature. The gene ADD1 is known to be associated with salt-sensitive essential hypertension–here we see the shared genetic architecture of that gene’s variants (represented as unique rsID accession identifiers) across the continuous cardiometabolic spectrum from health (systolic and diastolic BP) to disease (hypertension), facilitating the understanding of CVD trait genetics underlying health-to-disease transitions and endophenotypes. HeartBioPortal’s frontend design philosophy has always been to keep the user interface clean and minimalistic to ensure viewer comprehension and avoid potential confusion; therefore, additional details can be acquired from the hyperlinked data download, which is presented in a tidy .csv format amenable to follow-up analyses.

## 3 Methods

HeartBioPortal2.0 is comprised of >200 KLOCs written in a mixture of R, Python-3.8, JavaScript and Rust. The frontend framework is Vue.js and the backend is implemented in a combination of Django, Aerospike, ElasticSearch and PostgreSQL. HeartBioPortal2.0 sources and syncs its CVD-relevant genomics data from biological databases such as Ensembl, ClinVar, NHGRI-EBI GWAS catalog, gwasATLAS, OMIM, dbGaP, GTEx, CREEDS, HapMap, 1000Genomes, gnomAD, GEO, ArrayExpress, and OmicsDI. It also contains CVD-relevant data from large-scale GWAS consortiums such as CARDIoGRAMplusC4D, TOPMed, FinnGen, AFGen, MESA, MEGASTROKE, UK Biobank, CHARGE, Biobank Japan, MyCode, among others (including individual publications in top-tier journals that make GWAS datasets and summary statistics publicly available for secondary analysis). This includes highly specialized GWAS studies for CVD phenotypes such as, for example, tetralogy of fallot, which can be found under the congenital heart disease phenotype category in HeartBioPortal2.0.

### Genetic Association

GWAS summary statistics for cardiovascular and renal/metabolic related (e.g,. kidney function) traits were downloaded and filtered down for markers with reported p<=0.05. We did not attempt to further harmonize the p-values from different studies but present them as downloaded from each source. Our p-value filtering is intended to remove most markers with no phenotype-association signal but is not intended to signal genome-wide significance, which typically uses a much more stringent threshold. Users looking to see only the most significant association results (e.g. p<1E-5 or p<1E-8) should consider using the p-value slider on the HeartBioPortal2.0 gene pages. To annotate the filtered genetic markers, we queried myvariant.info (17) via post requests, and parsed the returned status and JSON using a custom Python script. Where possible, we attempted to use rsIDs from the original GWAS study to annotate each marker of interest. Where this was not possible, we queried using the chromosomal coordinates and Ref/Alt alleles (genomic HGVS). Returned annotated results for multi-allelic markers were split into the child bi-allelic markers, each keeping the original p-value for convenience. We chose to primarily rely on SnpEff (18) annotations because these were the most complete in terms of number of variants covered for the most relevant annotation fields presented in the data downloads.

In contrast to disease states which have a “yes” or “no” value for each patient, quantitative traits involve a continuous variable for each patient. For example, hypertension is a disease state while systolic blood pressure is a quantitative trait. Instead of the p-value indicating the association of a variant with risk of disease, quantitative GWAS studies report the association of a variant with a raised or lowered clinical value such as systolic blood pressure or some other trait. We have processed quantitative trait GWAS summary statistics the same as for disease state summary statistics. Both data types are present in new HeartBioPortal2.0 visualizations such as the variant viewer lollipop plot (Figure 2).

**Figure 2.**
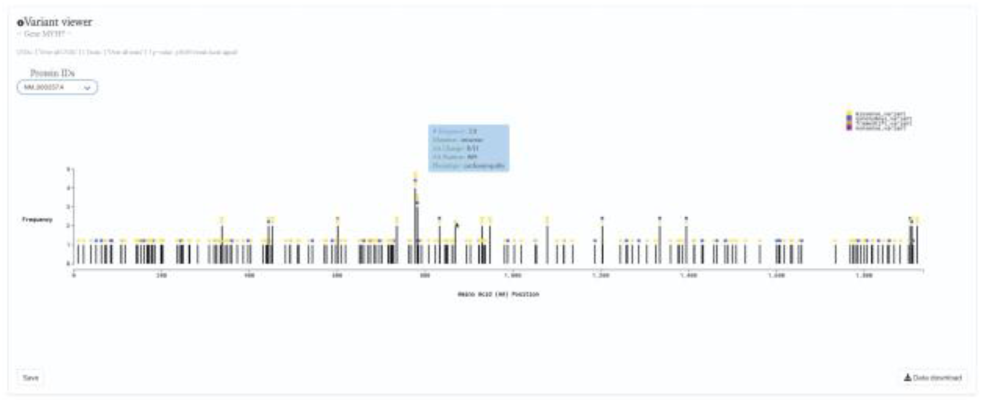
HeartBioPortal2.0’s variant viewer chart feature. The gene MYH7 is known to be associated with cardiomyopathy–here we see an interactive lollipop plot showing all known clinical genetic variants in MYH7 across multiple biological databases such as ClinVar, NHGRI-EBI GWAS Catalog, Ensembl, gnomAD, etc. The user can learn more information about any variant by hovering over the circles to display, e.g., the MYH7:p.Arg869His variant in this newly updated and sleek variant viewer chart. In addition, HeartBioPortal2.0 users can save their chart (for figures/diagrams in publications, seminars, etc.) and press the data download button to obtain additional details/metadata presented in a tidy .csv format amenable to additional follow-up data analyses. There are often multiple accession identifiers present in the dropdown menu of the variant viewer plot (e.g., for the gene DSP or TTN) to display variants found in both the canonical transcript of the gene as well as its alternative splice isoforms. Therefore, the variant viewer plot depicts variants found in all transcripts that have a known protein product associated with the respective CVD phenotype.

### Gene Expression

HeartBioPortal2.0 uses the GEO2R tool from Gene Expression Omnibus (GEO) (19–20), which relies on the limma (21) Bioconductor package to identify genes that are differentially expressed across experimental conditions (e.g., case/control), and applies a Benjamini & Hochberg (False discovery rate) (22) adjustment to the p-values, reporting only those top differentially expressed genes whose adjusted p-values are less than 0.05. Therefore, the expression results presented in HeartBioPortal2.0 are of high stringency to minimize the presence of any false positives.

Through the HeartBioPortal2.0 data processing pipeline, initial parsing of GEO for studies associated with CVD keywords returned a list of 685 CVD studies hosted on the GEO platform. Removal and manual curation of duplicate studies due to presence of multiple CVD keywords was performed, resulting in 541 unique CVD studies. Due to diversity in experimental type and study designs, manual curation of the remaining studies was performed to: 1) verify the type of experimental data, 2) verify that the research question was appropriate for the associated CVD keyword, and 3) verify proper selection of case/control subjects for differential gene expression analysis with respect to the CVD phenotype of interest.

This curation procedure ensured that the 142 remaining studies were appropriate for the gene expression feature of the HeartBioPortal2.0 platform, and these studies were input into the HeartBioPortal2.0 data processing pipeline. This pipeline utilizes the GEOquery (23) and limma packages in R through the GEO2R webtool to download experiment results and perform differential expression analysis and includes downstream processing to ensure a uniform standardized format. Differential expression analysis resulted in 60 carefully selected and curated studies with 9,641 more significant differentially expressed genes in HeartBioPortal2.0’s gene expression suite.

### System Performance and Security

Using the Rust programming language in HeartBioPortal2.0, we experienced 20X speedups (relative to Python in HeartBioPortal1.0) in creating a customized NoSQL database for our users to filter/download hundreds of gigabytes of genetic variant data in less than a few seconds. For example, registered users experience blazingly fast data downloads (<1s, 10^^^7 variants). Therefore, the Data Download button now has much faster performance across all major browsers (Firefox, Chrome, and Safari), in addition to being protected by reCAPTCHA and requiring user login/registration for authentication credentials. The NIH Genomic Data Sharing policy encourages minimal barriers to access genomic summary results. Therefore, investigators are able to see and query data at the summary level within HeartBioPortal2.0 without needing to go through a data access request process, since the data is de-identified. However, logging in via an authorized account is required before a user can download the de-identified data. Similar to other resources like dbGaP, this enables logging and monitoring of users and provides an opportunity to remind users of the three principles for responsible research use and compliance (no attempt to re-identify, use only for research or health purposes, and review of the responsible genomic data use information materials). The data provided in HeartBioPortal2.0 are available under the ODC Open Database License (ODbL). Users are free to share and modify the data so long as they properly attribute any public use of the database, or works produced from the database; keep the resulting datasets open; and offer their shared or adapted version of the dataset under the same ODbL license.

## 4 Discussion

Future directions in HeartBioPortal’s software development (in version 3.0 and above) will include further refining and updating complex visualizations like the genetic ancestry and variant geography maps (Figure 3), as well as including new bioinformatics pipelines and frontend support for harmonizing and visualizing CVD-relevant single cell RNA sequencing (scRNA-seq) and ATAC sequencing (scATAC-seq) datasets from multiple studies. Furthermore, to augment the wealth of CVD-relevant single nucleotide polymorphism (SNP) data from various GWAS efforts, we plan to add CVD-relevant structural variation (copy number variants, complex short substitutions, inversions, mobile elements, translocations, etc.), including drug discovery support (early target discovery genetics) to facilitate data-driven systematic target identification and prioritization of CVD drugs, effectively expanding the HeartBioPortal platform into a digital drug discovery dashboard for cardiovascular/renal/metabolic disease. In addition, we have constructed a product roadmap to interoperate HeartBioPortal with controlled-access data commons such as NHLBI’s BioData Catalyst (BDC), leveraging open-source tools such as Fence (24) to provide downstream authentication and authorization of Gen3 services (25), with the end-goal of transforming HeartBioPortal into a Gen3 client of BDC. This will give BDC users access to uniformly processed CRAM/VCF file results from a wide range of available human studies present in HeartBioPortal. Similar to our project, an existing resource called the eQTL Catalogue (26) processes public datasets using its uniform computational pipeline and we aim to create a similar sort of dynamic for interoperating BDC with HeartBioPortal using uniform bioinformatics pipelines developed and deployed securely at scale in the cloud. These computational innovations will create substantial real-world impact by introducing a novel hybrid computing strategy that turns a public database user login (e.g., from HeartBioPortal’s authentication page) into a secure key for unlocking additional controlled-access datasets in a safe and secure manner within the same UI (e.g., to see gene or variant-level results from data ecosystems like BDC, for users that have been pre-approved by BDC via a formal data access request made to an NHLBI data access committee to view specific datasets/cohorts).

**Figure 3.**
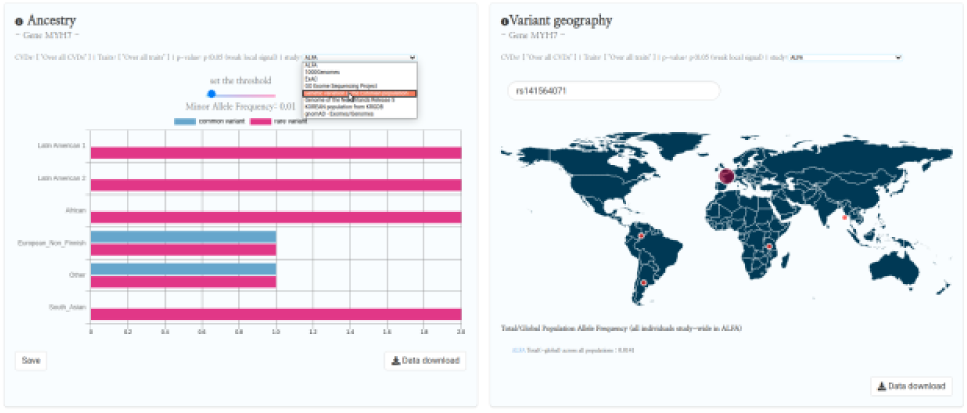
HeartBioPortal2.0’s genetic ancestry and variant geography feature. CVD-relevant exome/genome population allele frequency data displayed in the Ancestry and Variant Geography maps from dozens of studies/cohorts ranging from gnomAD to “Genome of the Netherlands Release 5” to “Korean population from KRGDB” among many others. In total, 54 diverse subpopulations across 27 new studies/cohorts have been added, relative to gnomAD alone in HeartBioPortal’s initial debut in May 2019. Users can now search individual mutations (rsID accession identifiers) as well as toggle a Minor Allele Frequency (MAF) slider bar that displays the proportion of rare vs. common variants at user-selected MAF thresholds.

## 5 Conclusions

HeartBioPortal2.0 constitutes a major upgrade to HeartBioPortal, both data-wise and features-wise, from frontend to backend. Plans are set in place to continue advancing the project rapidly along multiple development fronts, with the goal of harmonizing and integrating large-scale CVD genomics datasets that drive analytical insight on the complex biology of cardiovascular, cardiorenal, and cardiometabolic disorders.

## 6 Acknowledgements

BBK would like to thank Andrey Rzhetsky, Ph.D. for financial support of HeartBioPortal’s DigitalOcean cloud hosting servers from May 2019 to March 2020. SP would like to thank Yoonseo Lee and Maria Krunic for technical support and useful discussions. The authors declare no competing financial interests.

